# Inhibition of the sodium-dependent HCO_3_^-^ transporter SLC4A4, produces a cystic fibrosis-like airway disease phenotype

**DOI:** 10.1101/2021.12.14.472535

**Authors:** Vinciane Saint-Criq, Anita Guequén, Amber Philp, Sandra Villanueva, Tábata Apablaza, Ignacio Fernández-Moncada, Agustín Mansilla, Livia Delpiano, Iván Ruminot, Cristian Carrasco, Michael A. Gray, Carlos A. Flores

## Abstract

Bicarbonate secretion is a fundamental process involved in maintaining acid-base homeostasis. Disruption of bicarbonate entry into airway lumen, as has been observed in cystic fibrosis, produces several defects in lung function due to thick mucus accumulation. Bicarbonate is critical for correct mucin deployment and there is increasing interest in understanding its role in airway physiology, particularly in the initiation of lung disease in children affected by cystic fibrosis, in the absence of detectable bacterial infection. The current model of anion secretion in mammalian airways consists of CFTR and TMEM16A as apical anion exit channels, with limited capacity for bicarbonate transport compared to chloride. However, both channels can couple to SLC26A4 anion exchanger to maximise bicarbonate secretion. Nevertheless, current models lack any details about the identity of the basolateral protein(s) responsible for bicarbonate uptake into airway epithelial cells. We report herein that the electrogenic, sodium-dependent, bicarbonate cotransporter, SLC4A4, is expressed in the basolateral membrane of human and mouse airways, and that it’s pharmacological inhibition or genetic silencing reduces bicarbonate secretion. In fully differentiated primary human airway cells, SLC4A4 inhibition induced an acidification of the airways surface liquid and markedly reduced the capacity of cells to recover from an acid load. Studies in the *Slc4a4*-null mice revealed a previously unreported lung phenotype, characterized by mucus accumulation and reduced mucociliary clearance. Collectively, our results demonstrate that the reduction of SLC4A4 function induced a CF-like phenotype, even when chloride secretion remained intact, highlighting the important role SLC4A4 plays in bicarbonate secretion and mammalian airway function.

## INTRODUCTION

Bicarbonate (HCO_3_^-^) and chloride (Cl^-^) are actively secreted in the lumen of airways by the lining epithelial cells. Even though, the pathophysiological consequences of decreased secretion of these anions has been extensively documented in cystic fibrosis (CF), the most common, autosomal recessive, disease in humans, there is still much discussion whether HCO_3_^-^ secretion *per se* affects airway homeostasis (1, 2). Impaired Cl^-^ secretion reduces the volume of the fluid that covers the airway epithelium, the airway surface liquid (ASL), leading to ciliary dysfunction and promoting mucus stasis and airway obstruction (3, 4). Deficient HCO_3_^-^ secretion reduces ASL pH which compromises post-secretory mucin maturation and clearance (5), impairs the antimicrobial function of epithelial cells (6, 7) and increases fluid absorption, further decreasing ASL hydration (8). Whilst, direct measurement of pH in the distal airways of CF children showed no acidification of the ASL (9), it has been demonstrated that the addition of HCO_3_^-^ induced an increase in ASL height in human airway epithelial cells (hAECs) cultures and restored the normal properties of mucus from CF patients (8, 10). Moreover, the use of aerosolized HCO_3_^-^ into CF-pig airways increased bacterial killing, clearly indicating that HCO_3_^-^ supplementation can correct inherent defects of CF in the airways (6, 11).

Even though, impaired HCO_3_^-^ secretion is recognised as a detrimental component of CF airway disease, a full mechanistic understanding of the process and players involved in transcellular HCO_3_^-^ transport in the airways is lacking. In normal airways, HCO_3_^-^ is secreted by CFTR and TMEM16A channels, but after inflammatory signalling HCO_3_^-^ secretion is augmented through increased expression of the Cl^-^ /HCO_3_^-^ exchanger Pendrin (SLC26A4) (12–16). Importantly, the mechanism of basolateral HCO_3_^-^ transport/uptake in native pulmonary tissues is still lacking. We reasoned that the identification and functional inhibition of such basolateral membrane proteins could be used as a proof-of-concept to better understand the role of HCO_3_^-^ secretion in airway homeostasis without altering Cl^-^ secretion. Here using *in vitro* fully differentiated human bronchial epithelial cells, we identified the Na^+^-dependent, HCO_3_^-^ electrogenic cotransporter (NBCe1), SLC4A4, as the basolateral protein responsible for HCO_3_^-^ influx, which couples to apical CFTR for HCO_3_^-^ secretion into the ASL. Importantly, SLC4A4 inhibition induced acidification of ASL, revealing the pivotal role of this cotransporter in airway pH homeostasis. These observations were further tested in murine models, and revealed that SLC4A4 is involved in both basal and Ca^2+^-stimulated HCO_3_^-^ secretion in mice airways. Strikingly, an *Slc4a4*^-/-^ mouse model showed significant pathological signs of muco-obstructive disease and reduced mucociliary clearance, confirming that inhibition of HCO_3_^-^ secretion alters airway homeostasis, mimicking what has been observed in human CF, and identifying a critical role played by SLC4A4.

## METHODS

### *In vitro* Human airway epithelial cells studies

#### Cell culture

Primary hAECs were a kind gift from Dr. Scott H. Randell (Marsico Lung Institute, The University of North Carolina at Chapel Hill, United States). The cells were obtained under protocol #03-1396 approved by the University of North Carolina at Chapel Hill Biomedical Institutional Review Board. Cells were expanded using the conditionally reprogrammed cell (CRC) culture method as previously described (Suprynowicz et al., 2012). Briefly, cells were seeded on 3T3J2 fibroblasts inactivated with mitomycin C (4 mg/ml, 2 h, 37°C) and grown in medium containing the ROCK inhibitor Y-27632 (10 mM, Tocris Biotechne, #1254) until they reached 80% confluence. Cells then underwent double trypsinization to remove the fibroblasts first and then detach the hAECs from the P150 dish. At that stage, cells were counted and could be frozen down. Cryopreserved cells were seeded onto semi-permeable supports (6.5 or 12 mm) in bilateral differentiating medium (ALI medium) as previously described (Randell et al., 2011). The apical medium was removed after 3-4 days and cells were then allowed to differentiate under air-liquid interface (ALI) conditions. Ciliogenesis started approximately 12–15 days after seeding and cells were used for experiments between days 25 and 35 after seeding.

#### Intracellular pH measurements

Primary airway epithelial cells, grown on 12 mm Transwell inserts, were loaded with the pH-sensitive, fluorescent dye BCECF-AM (10μM, ThemoFisher Scientific #B-1150) for 1h in a Na-HEPES buffered solution (130 mM NaCl, 5 mM KCl, 1 mM MgCl_2_, 1 mM CaCl_2_, 10 mM Na-HEPES, and 5 mM D-glucose set to pH 7.4) at 37°C. Cells were mounted on to the stage of a Nikon fluor inverted microscope and perfused with a modified Krebs (KRB) solution (115 mM NaCl, 5 mM KCl, 25 mM NaHCO_3_, 1 mM MgCl_2_, 1 mM CaCl_2_, and 5 mM D-glucose) gassed with 5% (v/v) CO_2_/95% (v/v) O_2_ or with a Na-HEPES-buffered solution gassed with 100% O_2_. Solutions were perfused across the apical and basolateral membranes at 37°C at a speed of 3 ml min^−1^ and 6 ml min^−1^, respectively. To test the sodium dependence of bicarbonate transport, a Na^+^-free KRB solution was used in which 115 mM NMDG-Cl replaced NaCl, and 25 mM choline-HCO_3_ replaced NaHCO_3_. To measure the effect of NBC inhibition on the recovery from CO_2_-induced acidification, epithelial cells were perfused basolaterally with 100 μM DMA (dimethyl amiloride, Sigma-Aldrich #A4562) to inhibit sodium-dependent hydrogen exchangers (NHEs) and 30 μM S0859 (Sigma-Aldrich #SML0638) to inhibit NBC. Intracellular pH (pHi) was measured using a Life Sciences Microfluorimeter System in which cells were alternately excited at 490 and 440 nm wavelengths every 1.024 s with emitted light collected at 510 nm. The ratio of 490 to 440 nm emission was recorded using PhoCal 1.6 b software and calibrated to pHi using the high K^+^/nigericin technique (17) in which cells were exposed to high K^+^ solutions containing 10 μM nigericin, set to a desired pH, ranging from 6 to 7.5. For analysis of pHi measurements, ΔpHi was determined by calculating the mean pHi over 60 s resulting from treatment. The initial rate of pHi change (ΔpHi/Δt) was determined by performing a linear regression over a period of at least 40 s.

#### Short-Circuit Current Measurements in human airway epithelial cells

Cells grown on 6.5 mm inserts were mounted into the EasyMount Ussing Chamber Systems (VCC MC8 Physiologic Instrument, tissue slider P2302T) and bathed in bilateral Cl^-^ free HCO_3_^-^ KRB (25 mM NaHCO_3_, 115 mM Nagluconate, 2.5 mM K_2_SO_4_, 6.0 mM Ca-gluconate, 1 mM Mg-gluconate, 5 mM D-glucose) continuously gassed and stirred with 5% (v/v) CO_2_/95% (v/v) O_2_ at 37°C or in bilateral NaHEPES buffered solution continuously gassed and stirred with 100% O_2_ at 37°C. Monolayers were voltage-clamped to 0 mV and monitored for changes in short-circuit current (ΔI_sc_) using Ag/AgCl reference electrodes. The transepithelial short-circuit current (I_sc_) and the Transepithelial electrical resistance (R_te_, expressed in Ω cm^2^) were recorded using Ag–AgCl electrodes in 3 M KCl agar bridges, as previously described (18), and the Acquire & Analyze software (Physiologic Instruments) used to perform the analysis. Cells were left to equilibrate for a further 10 min before amiloride (10 μM, apical, Sigma-Aldrich #A7410) and S0859 (30 μM, basolateral) were added. Results were normalized to an area of 1 cm^2^ and expressed as Isc (μAmp.cm^-2^). The number of replicates was determined using previously obtained short circuit current measurements.

#### ASL pH Measurements

ASL pH measurements were performed as previously described (19, 20). Briefly, cells grown on 6.5 mm transwells were washed apically with modified Krebs solution for 15 min at 37°C, 5% CO_2_. The ASL was stained using 3 μl of a mixture of dextran-coupled pH-sensitive pHrodo Red (0.5 mg/ml, λex: 565 nm, λem: 585 nm; ThermoFisher Scientific, #P10361) and Alexa Fluor® 488 (0.5 mg/ml, λex: 495 nm, λem: 519 nm; ThermoFisher Scientific #D-22910) diluted in glucose-free modified Krebs buffer, overnight at 37°C, 5% CO_2_. The next day, fluorescence was recorded using a temperature and CO_2_-controlled plate reader (TECAN SPARK 10M) and forskolin (Tocris Biotechne #1099) and S0859 were added basolaterally at indicated times. The ratio of pHrodo to Alexa Fluor® 488 was converted to pH using a calibration curve obtained by clamping apical and basolateral pH in situ using highly buffered solutions between 5.5 and 8 (19). To prevent inter-experiment variability, the standard curve calibration was performed on each independent experiment.

#### RNA extraction and PCR analysis

RNA isolation from cells was performed using PureLink® RNA Mini Kit (Ambion, Life technologies, #12183018A), following the manufacturer’s instructions. Briefly, lysates were mixed with 70 % ethanol and loaded onto a silica-membrane column. Columns were washed with different buffers and total RNA was eluted in DNAse and RNAse-free water and stored at -80°C until use. DNase treatment was performed on 300 ng RNA prior to Reverse Transcription Polymerase Chain Reaction (RT-PCR) using RNAse-free DNAse I (Roche, # 04716728001) at 37°C for 10 min. Reaction was then stopped by increasing the temperature to 70°C for 10 min. Complementary DNA (cDNA) was synthesized from total RNA (300 ng) using M-MLV Reverse Transcriptase (Promega, #M1701) as per supplier’s protocol (1hr at 37°C followed by 10 min at 70°C). Expression of SLC4A family members was evaluated by PCR using specific primers (Sup Table 1), in a total volume of 25 μL containing 2 μL cDNA template, 5 μL 5X Q5 reaction buffer, 0.5 μL 10 mM dNTP, 1.25 μL 10 μM of each primer, 0.25 μL Q5 high Fidelity Polymerase (New England Biolabs Inc., M0491), 5 μL 5X Q5 high enhancer (Denaturation, 98°C, 30 seco; (98°C 5 sec, 72°C 30 sec -72°C 20 sec) x35 cycles; Final Extension, 72°C, 2 min. PCR products were loaded onto a SYBR Safe DNA stain (Life Technologies, cat. no. S33102)-containing, 2 % agarose gels in TBE and electrophoresis was ran at 90V for 1.5hr. Amplified products were visualized using LAS-3000 Imaging System (Fuji).

### *Ex vivo* murine airway studies

#### Animals

The *Slc4a4*^-/-^ mice was obtained from its laboratory of origin(21) and bred in the original 129S6/SvEv/Black Swiss background or C57BL/6J. As observed in Fig S4A animals maintained in the C57BL/6J background were severely affected by lethality before weaning (day 21 post birth). Therefore experiments were performed in the hybrid animals. The wild type C57BL/6J mice were from The Jackson Laboratories (USA). Animals were bred and maintained in the Specific Pathogen Free mouse facility of Centro de Estudios Científicos (CECs) with access to food and water ad libitum. 8-12 weeks-old or 16-20 days-old, male or female mice were used. Unless otherwise stated, all procedures were performed after mice were deeply anesthetized via i.p. injection of 120 mg/kg ketamine and 16 mg/kg xylazine followed by exsanguination. All experimental procedures were approved by the Centro de Estudios Científicos (CECs) Institutional Animal Care and Use Committee (#2015-02) and are in accordance with relevant guidelines and regulations.

#### Ussing chamber experiments

Tracheae were placed in P2306B of 0.057 cm^2^ (Fig 3A-K and Fig S4A-F) or P2307 of 0.031 cm^2^ (Fig 5E) tissue holders and placed in Ussing chambers (Physiologic Instruments Inc., San Diego, CA, USA). Tissues were bathed with bicarbonate-buffered solution (pH 7.4) of the following composition (in mM): 120 NaCl, 25 NaHCO_3_, 3.3 KH_2_PO_4_, 0.8 K_2_HPO_4_, 1.2 MgCl_2_, 1.2 CaCl_2_ or HEPES buffer: 130 NaCl, 5 KCl, 1 MgCl2, 1 CaCl2, 10 Na-HEPES (pH adjusted to 7.4 using HCl); supplemented with 10 D-Glucose, gassed with 5% CO_2_–95% O_2_ (bicarbonate buffer) or 100% O_2_ (HEPES buffer) and kept at 37°C. The transepithelial potential difference referred to the serosal side was measured using a VCC MC2 amplifier (Physiologic Instruments Inc.). The short-circuit currents were calculated using the Ohm’s law as previously described(22). Briefly electrogenic Na^+^absorption was inhibited using 10 μM amiloride (Sigma #A7410), cAMP-dependent anion secretion was induced using an IBMX+Forskolin mixture of 100 μM IBMX (Sigma #I5879) + 1 μM forskolin (Sigma #F6886), Ca^2+^-dependent anion secretion was induced by 100 μM UTP (Sigma #U6750). Acetazolamide (100 μM) was used to inhibit bicarbonate production (Sigma #A7011), to block SLC4A4 30 μM S0859 was used (kindly donated by Juergen Puenter, Sanofi-Aventis, France and dissolved in ethanol), and 30 μM CACC_inh_A01 (Calbiochem) to inhibit CFTR and TMEM16A channels. The ΔI_sc_ values were calculated by subtracting I_sc_ values values before from I_sc_ values after the addition of drugs but UTP induced current was calculated as the area under the curve (A.U.C.) of the first 5 minutes post UTP addition using the Acquire & Analyze 2.3v software. Tissues with R_te_ values below 50 Ωcm^2^ were discarded as were not suitable for *bona fide* electrophysiological determinations (23).

#### Airway cells isolation and intracellular pH determinations

Tracheae were incubated with Pronase 25 μg/ml at 37°C for 30 minutes. Then the trachea was placed in a petri dish with DMEM-F12, and the airway epithelium was dissociated by scrapping with tweezers, the cells were collected and spun at 3000 r.p.m. for 5 minutes at room temperature and the supernatant was removed. The cell pellet was incubated with 500 μl of trypsin 1X at 37°C for 5 minutes and centrifuged at 3000 r.p.m for 5 minutes at room temperature and the supernatant was removed. The airway epithelial cells were resuspended into 150 μl of DMEM-F12 medium supplemented with 10% fetal bovine serum (FBS) and seeded on poly-lysine coated 25 mm glass coverslips in 35 mm Petri dishes. Freshly isolated cells from mouse trachea were loaded with 0.5 uM BCECF-AM (ThermoFisher Scientific #B-1170) for 10 min at 37°C. After loading, cells were washed and incubated 30 min in imaging solution (see below) to allow probe de-esterification. Cells were mounted into an open chamber and superfused with a bicarbonate buffer imaging solution of the following composition (in mM): 120 NaCl, 25 NaHCO_3_, 3.3 KH_2_PO_4_, 0.8 K_2_HPO_4_, 1.2 MgCl_2_, 1.2 CaCl_2_ and bubbled with air/5% CO_2_. For experiments without bicarbonate, same HEPES buffer as in Ussing chamber experiments was added to the imaging buffer and bubbled with 100% O_2_. In the low chloride bicarbonate buffer, 120 mM NaCl was replaced by 115 mM 115 mM Na-Gluconate and 5 mM NaCl. Experiments were carried out on an Olympus IX70 inverted microscope equipped with a 40X oil-immersion objective (NA 1.3), a monochromator (Cairn, UK) and a CCD Hamamatsu Orca camera (Hamamatsu, Japan), controlled by Kinetics software. All solutions were superfused at 37°C using an in-line heating system (Warner instruments). BCECF was excited sequentially at 490 nm and 440 nm for 0.05-0.1 s and emission collected at 535/15 nm. The F490/F440 ratio was computed and transformed to pH units by performing a pH-clamp. Briefly, cells were exposed to 5 μM nigericin and 20 μg/ml gramicidin in a buffer composed of (in mM) 10 HEPES, 129 KCl, 10 NaCl, 1.25 MgCl2, 1 EGTA, 10 glucose, with pH values ranging between 6.8-7.8 and the observed changes in fluorescence were quantified and used to construct a calibration curve.

#### Airway cell isolation and RT-PCR

Tracheae were incubated with Pronase 25 μg/ml at 37°C for 30 minutes. Trachea was placed in DMEM-F12 with 10 mM D-glucose and epithelium was isolated by scrapping with tweezers and further homogenized in 250 μl Trizol (Trizol Reagent) and RNA isolated following the manufacturer’
ss instructions. The dried pellet of RNA was resuspended with 35 μl nuclease free water and stored at -80°C. Genomic DNA was removed through DNase treatment. The concentration and integrity of RNA isolation was verified using a NanoDrop spectrophotometer Maestrogen. RNA was reverse transcribed into cDNA using the ImProm-IITM Reverse Transcription System (Promega) following manufacturer’s recommendations. The specific primer pairs used for *Slc4a4, Slc4a5, Slc4a7, Slc4a8, Slc4a10, Slc4a4*-A and *Slc4a4*-B PCR amplification are provided in Sup Table 2. PCR amplification was performed starting with 3-minute template denaturation step at 95°C, followed by 40 cycles of denaturation at 95°C for 30 seconds and combined primer annealing/extension at temperature as appropriate.

#### Histology and immuno fluorescence

Human tissues were obtained from the Pathological Anatomy Subdepartment of Hospital Base Valdivia (Valdivia, Chile) and corresponded to surgical resections of lung tumors that contained normal parenchyma including epithelium. The studies were approved by the Bioethical Committee for Human Research of the Servicio de Salud Los Ríos (Submitted). To obtain mice tissues the animals were placed in a 1-litre induction chamber under 1000 ml min^−1^ flow of air containing 2.5% isoflurane. Then kept under anaesthesia with 2% isoflurane at a constant flow rate of 500 ml min^−1^ using a mask. Mice were euthanized by exsanguination by severing the inferior vena cava under deep anaesthesia. Mice were perfused with 4% paraformaldehyde (PFA). Trachea and lung were removed and incubated overnight in 4% PFA at 4°C. Paraffin sections (4 μm) were treated with Trilogy ™ 1X (Cell Marque cat# 920P-06), blocked with 2.5% normal goat serum (Vector Laboratories cat# S-1012), and incubated with 1:100 anti-NBCe1 (anti SLC4A4; Alomone cat#ANT-075) 4°C overnight. Sections were incubated with secondary antibody 1:2,000 anti-rabbit Alexa Fluor 488 (Invitrogen cat# A-11008) 2h at room temperature. For colocalization 1:100 anti-NBCe1 was incubated with 1:1,000 anti-Clara Cell Secretory Protein (CCSP; Merck Millipore cat#07-623) or 1:200 anti-alpha tubulin (Santa Cruz cat#sc-5286) overnight at 4°C, and incubated with secondary antibody 1:2,000 anti-rabbit Alexa Fluor 568 (Life Technologies cat#A-11011 or 1:2,000 anti-mouse Alexa Fluor 568 (Life Technologies cat#A-11004) respectively. Nuclei were stained with 1:2,000 propidium iodide (Invitrogen cat#P21493). All immunofluorescence images were captured using a confocal microscope (Olympus FV1000).

#### Whole-mount trachea Alcian blue cartilage staining

Tissue was fixed in 95% ethanol overnight followed by 2h staining with 0.03% Alcian blue (Sigma cat#A5268) dissolved in 80% ethanol and 20% acetic acid. Samples were cleared in 2% KOH, and pictures taken under a stereomicroscope.

#### Mucociliary clearance determination

Speed of polystyrene beads in trachea samples was determined as previously described(22). Briefly, the tracheas were isolated and mounted with insert needles onto extra thick blot paper (Bio-Rad) and transferred into a water-saturated chamber at 37°C. The filter paper was perfused with HCO_3_^-^ buffered solution of the following composition (in mM): 120 NaCl, 25 NaHCO_3_, 3.3 KH_2_PO_4_, 0.8 K_2_HPO_4_, 1.2 MgCl_2_, 1.2 CaCl_2_ (gassed with 5% (v/v) CO_2_/95% (v/v) O_2_ to maintain solution pH close to 7.4) at a rate of 1 ml/min and at 37°C. Polystyrene black dyed microspheres (diameter 6μm, 2.6% solid-latex, Polybead, Polyscience Inc) were washed and resuspended in HCO_3_^-^ or HEPES solution and 4 μl of particle solution with 0.3% latex were added onto the mucosal surface of the trachea. Particle transport was visualized every 5 seconds for 15 min using a ZEISS SteREO Discovery.V12, with digital camera Motic (Moticam 5.0). Particle speed was calculated with NIH ImageJ software and speed of MCC was expressed in μm/sec. To inhibit HCO_3_^-^ secretion, HCO_3_^-^ solution was substituted for the HCO_3_^-^ free solution (HEPES) of the following composition (in mM): 145 NaCl, 1.6 K_2_HPO_4_, 0.4 KH_2_PO_4_, 1.0 MgCl_2_, 1.3 CaCl_2_ throughout the experiments. In these experiments, the HCO_3_^-^ free solution (HEPES) was gassed with 100% O_2_ gas to maintain solution pH close to 7.4. Beads tracked on each tissue were averaged and were used as corresponding to one tissue sample.

### Statistical analysis

Experiments in human cells were analysed using GraphPad Prism v9. Statistical analysis was performed taking into account the number n of independent repetitions (done in different days) of the experiments (stated in the Figure legends) using cells at different passage numbers from 3 different donors. Thus n numbers given in the figure legends are considered as biological replicates. Statistical tests used are indicated in the figure legends and p-values are shown in the figures. Experiments using animals were analysed using the Sigmaplot 12 software. Statistical analysis was performed considering as n the number of animals used as source for tissues. These n numbers given on figure legends are considered biological replicates. For pHi experiments isolated cells were from at least 3 different animals per group and statistical analysis was performed using values from individual cells. ANOVA on Ranks was used for comparisons of more than 2 data sets while Rank Sum test for comparison of data with 2 data sets. For survival analysis Log Rank test was used. P-value of <0.05 was considered statistically significant. Sample size for experiments in human cells were calculated using Cohen’s d, a power analysis showed that the sample size of 5 independent experiments has a 90% power to detect a difference, assuming a 5% significance level and a two-sided test. Calculations were based in previously published data for intracellular pH (17), ASL pH measurements (20) and unpublished data set for short-circuit currents. Sample size for animal experiments was calculated using previous published data for Ussing chamber experiments (22, 23), intracellular pH (24) and mucociliary clearance (22).

### Data availavility

All data analysed in this study is presented in the manuscript and in the supplementary data.

## RESULTS

### Primary human airway epithelial cells express bicarbonate transporters of the SLC4A family

We first investigated, by PCR, whether primary hAECs expressed different members of the SLC4A family of Na^+^-coupled HCO_3_^-^ transporters (NCBT) and isolated RNA from kidney and the Calu-3 cell line, which was derived from a metastatic site of a lung adenocarcinoma, as positive controls. The use of specific primers (Table S1) for each NCBT revealed that SLC4A4, A5 and A8 are expressed at mRNA levels in primary hAECs from 3 different individuals (P1, P2, P3, Fig S1A). SLC4A7 and SLC4A10 showed very low to none level of mRNA expression (Fig S1A). Interestingly, isoforms B/C of NBCe1 (known as the pancreatic isoform) were more highly expressed than isoform A (known as the kidney isoform).

### SLC4A4 is central for bicarbonate secretion and intracellular pH homeostasis in human airway cells

We then tested whether there was an active NCBT under resting conditions in primary hAECs. The cell cultures were mounted in Ussing chambers in buffers containing either HCO_3_^-^ (but no Cl^-^) or HEPES, and treated basolaterally with the inhibitor S0859 (30 μM). Results, shown in Figure 1 confirmed that, under basal conditions, HCO_3_^-^ secretion was inhibited by S0859 (Fig 1A,B) and that this pharmacological inhibitor did not have any effect on short-circuit current (Isc) in the absence of HCO_3_^-^ (Fig 1C,D). These results show that there is an electrogenic HCO_3_^-^ transporter at the basolateral membrane of hAECs, which is consistent with SLC4A4, since SLC4A7 and A8 are electroneutral, and SLC4A5 has been shown to be localized to the apical membrane of renal epithelial cells (25). Next, we used intracellular pH (pH_i_) measurements to functionally investigate NBCe1 activity using a CO_2_-induced acidification protocol (26). As shown in Figure 1E, exposing cells bilaterally to a HCO_3_^-^/CO_2_-gassed KRB solution induced a transient acidification. On the other hand, an apical-only CO_2_ exposure, in the absence of basolateral HCO_3_^-^, induced a sustained acidification (of the same amplitude as with bilateral HCO_3_^-^/CO_2_, Figure 1F,H), that recovered when HCO_3_^-^ was re-introduced basolaterally (Figure 1F,I). This pH_i_ recovery depended on the presence of Na^+^ in the basolateral solution (Figure 1G) consistent with a Na^+^-coupled HCO_3_^-^ transporter, which was confirmed using S0859 which blocked the pH_i_ recovery from the CO_2_-induced acidification. In order to isolate NBCe1 only changes in pH_i_, the contribution of Na^+^/H^+^ exchanger NHE was inhibited using 100 μM Dimethyl amiloride (DMA). In this condition, S0859 still significantly decreased the rate of pH_i_ recovery from the CO_2_-induced acidification (Figure 1J,K).

**Figure 1.**
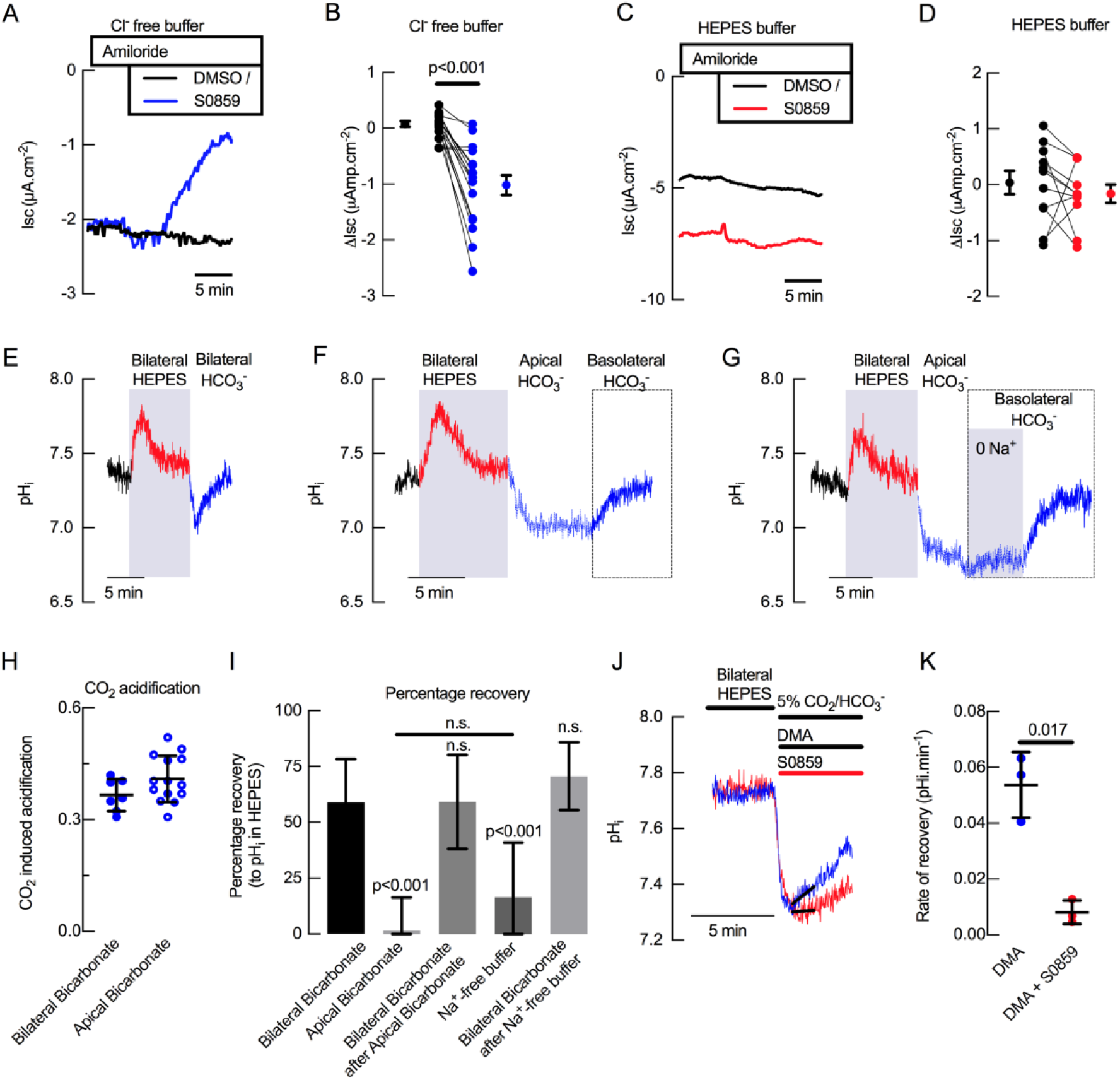
Basal bicarbonate secretion requires SLC4A4 activity in primary hAECs. Representative traces of Isc in (A) Cl^-^ free solution and (C) HCO_3_^-^ free solution of primary hAECs cultures. Summary of S0859-induced changes in I_sc_ in the presence (B) or absence (D) of HCO_3_^-^ in primary hAECs cultures (on panels B and D, each dot represents an independent experiment; means ± sem are shown next to the individual data; respectively n=17 and n=11 independent experiments using cells from N=3 donors; two-tailed, paired t-test). (E) Representative pH_i_ trace of primary hAECs bathed first in bilateral HCO_3_^-^ KRB (gassed with 5% CO_2_) then bilateral Hepes buffered KRB (no HCO_3_^-^, no CO_2_) and then bilateral HCO_3_^-^ KRB (gassed with 5% CO_2_). CO_2_ removal and re-introduction is marked by a transient increase and decrease in pH_i_ respectively. (F) Representative trace of pH_i_ recovery after CO_2_-induced acidification in the absence of basolateral HCO_3_^-^. (G) Representative trace of pH_i_ recovery after CO_2_-induced acidification in the absence of basolateral Na^+^ and HCO_3_^-^. (H) Summary of the magnitude of CO_2_ -induced acidification (bilateral bicarbonate, n=7, apical bicarbonate n=14, unpaired t-test, bars represent mean ± standard deviation (S.D.)). (I) Mean percentage of pH_i_ recovery after perfusion of the different solutions (Bilateral Bicarbonate, n=7; Apical Bicarbonate, n=14; Bilateral Bicarbonate after Apical Bicarbonate^-^, n=5; Na^+^-free buffer, n=8; Bilateral Bicarbonate after Na^+^-free buffer, n=8; One-way ANOVA with Holm-Sidak correction for multiple comparison tests, bar graph represents mean ± S.D.). (J) Representative trace of intracellular pH measurements showing recovery from CO_2_-induced acidification in the presence (red line) or absence (blue line) of NBC inhibitor S0859. (K) Summary of rates of recovery from CO_2_-induced acidification in the presence of NHE inhibitor (Dimethyl Amiloride, DMA, 100 μM) and in the presence (red bar) or absence (blue bar) of S0859 (30 μM), (n = 3, paired t-test; bars represent mean ± S.D.).

### Basolateral HCO_3_^-^ uptake is essential for ASL pH homeostasis

In order to test whether HCO_3_^-^ transport by basolateral SLC4A4 impacted apical HCO_3_^-^ secretion, we measured the effect of S0859 on ASL pH. First, S0859 was added basolaterally to primary hAECs and ASL pH continuously measured. NBCe1 inhibition significantly decreased ASL pH under resting conditions (Fig 2A,B), and partially prevented the forskolin-induced increase in ASL pH which we have previously shown was via CFTR (19, 20)(Fig 2C,D). Moreover, when S0859 was added after forskolin, it significantly reduced the forskolin-induced, CFTR-dependent, increase in ASL pH (Fig 2E,F) confirming the central role of SLC4A4 cotransporter in ASL pH homeostasis under both resting and stimulated conditions.

**Figure 2.**
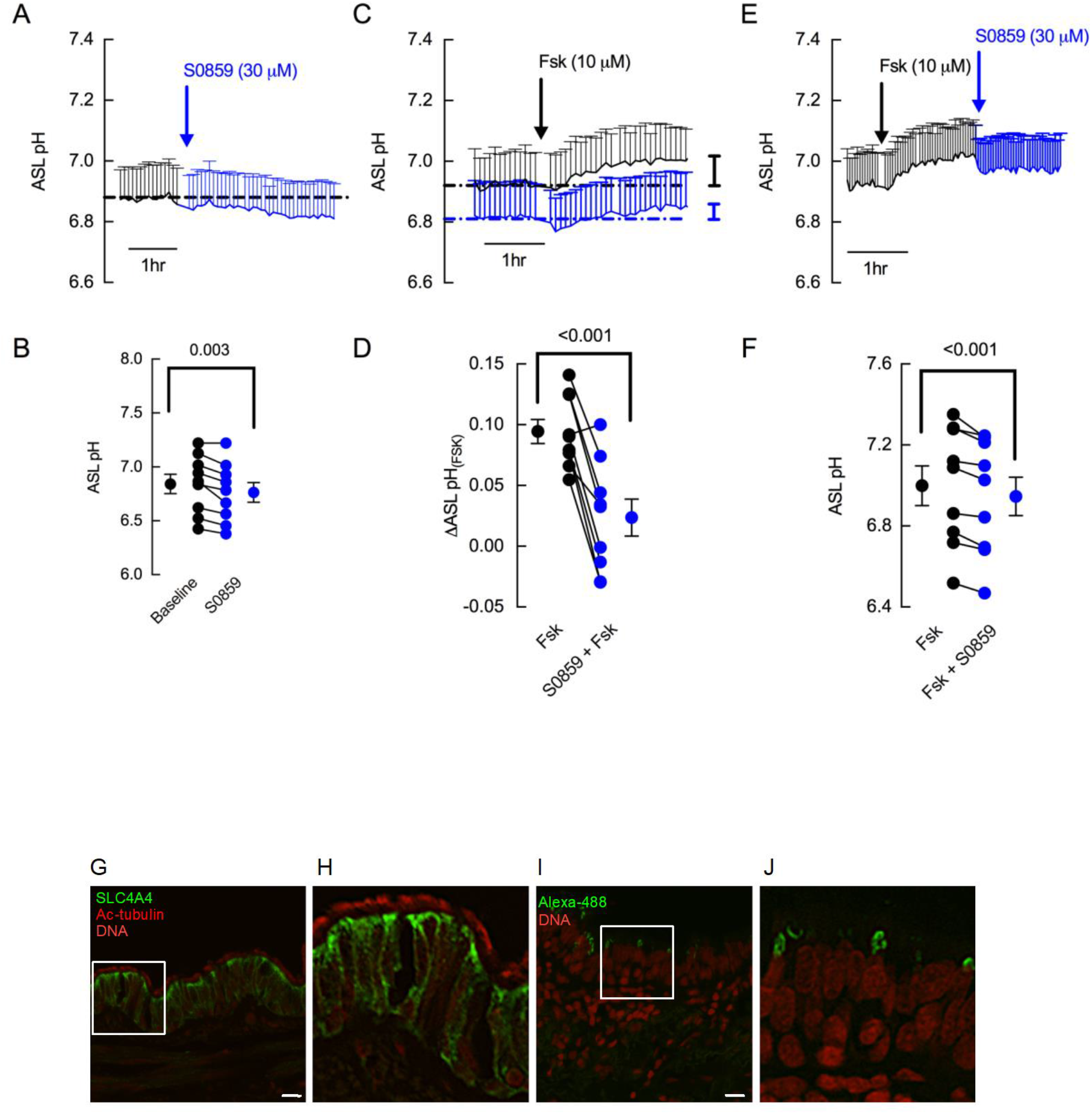
S0859 decreases resting ASL pH in primary human airway epithelial cells (hAECs), by blocking basolateral SLC4A4. (A) Mean (+S.E.M.) trace of ASL pH recordings. ASL pH was measured under resting conditions for 1.5hr before S0859 (30 uM) was added basolaterally. (B) Mean resting ASL pH before (black circles) and after (blue circles) addition of basolateral S0859 (n=9 independent experiments performed on epithelial cells from N=3 donors; paired t-test). (C) Mean (+SEM) traces of ASL pH from hAECs pre-treated for 3 h with vehicle control (DMSO, black trace) or S0859 (30 μM, basolateral, blue trace). (D) Mean Forskolin (Fsk)-induced changes in ASL pH in hAECs treated with DMSO (black circles) or S0859 (blue circles) (n=9 independent experiments performed on epithelial cells from N=3 donors; paired t-test). (E) Mean (+SEM) traces of ASL pH from hAECs treated with Fsk for 2.5 h and then S0859. (F) Mean Fsk-stimulated ASL pH before (black circles) and after (blue circles) addition of basolateral S0859 (n=9 independent experiments performed on epithelial cells from N=3 donors; paired t-test). (G-H) SLC4A4 locates in basolateral membrane of acetylated tubulin (Ac-tubulin) positive human airway epithelial cells. (I-J) correspond to negative controls for anti-SLC4A4 omitted antibody. Representative images of 3 different samples. Bar 20 μM.

Finally, immunolocalization of SLC4A4 protein in human airways showed intracellular and basolateral membrane staining in epithelial cells that were also positively stained for acetylated-tubulin indicating that SLC4A4 is preferentially expressed in ciliated cells in human airways (Fig 2G,H).

### Bicarbonate secretion is calcium-activated in mouse trachea

To investigate the expression of SLC4 exchangers in mouse airway epithelium, we performed RT-PCR of epithelial cells from mouse tracheas and observed that several members of the SLC4 family including *Slc4a4, Slc4a5, Slc4a7* and *Slc4a10* were expressed (Fig S2). Studies of *Slc4a4* isoforms demonstrated that isoform B/C but not isoform A was expressed in mouse airways. Next we characterized HCO_3_^-^ secretion in the mouse. Ussing chamber experiments performed in freshly excised mouse trachea using HCO_3_^-^ (Fig 3A) or HEPES (Fig 3B) buffered solutions, showed that UTP-induced an anion current that was significantly reduced in the absence of HCO_3_^-^ (−138 ± 28 vs -82 ± 15 μA cm^-2^; p<0.01 Mann-Whitney test), with no significant effect on the cAMP-induced anion secretion, or the amiloride-sensitive sodium absorption (Fig 3C). Complementary studies in HCO_3_^-^ buffer showed that *de novo* synthesis of HCO_3_^-^ was not participating in the UTP-induced electrogenic anion secretion, as incubation of tracheas with acetazolamide didn’t affect the magnitude of the UTP-induced current (−124 ± 14 μA cm^-2^; p>0.05 One-way ANOVA).

**FIGURE 3.**
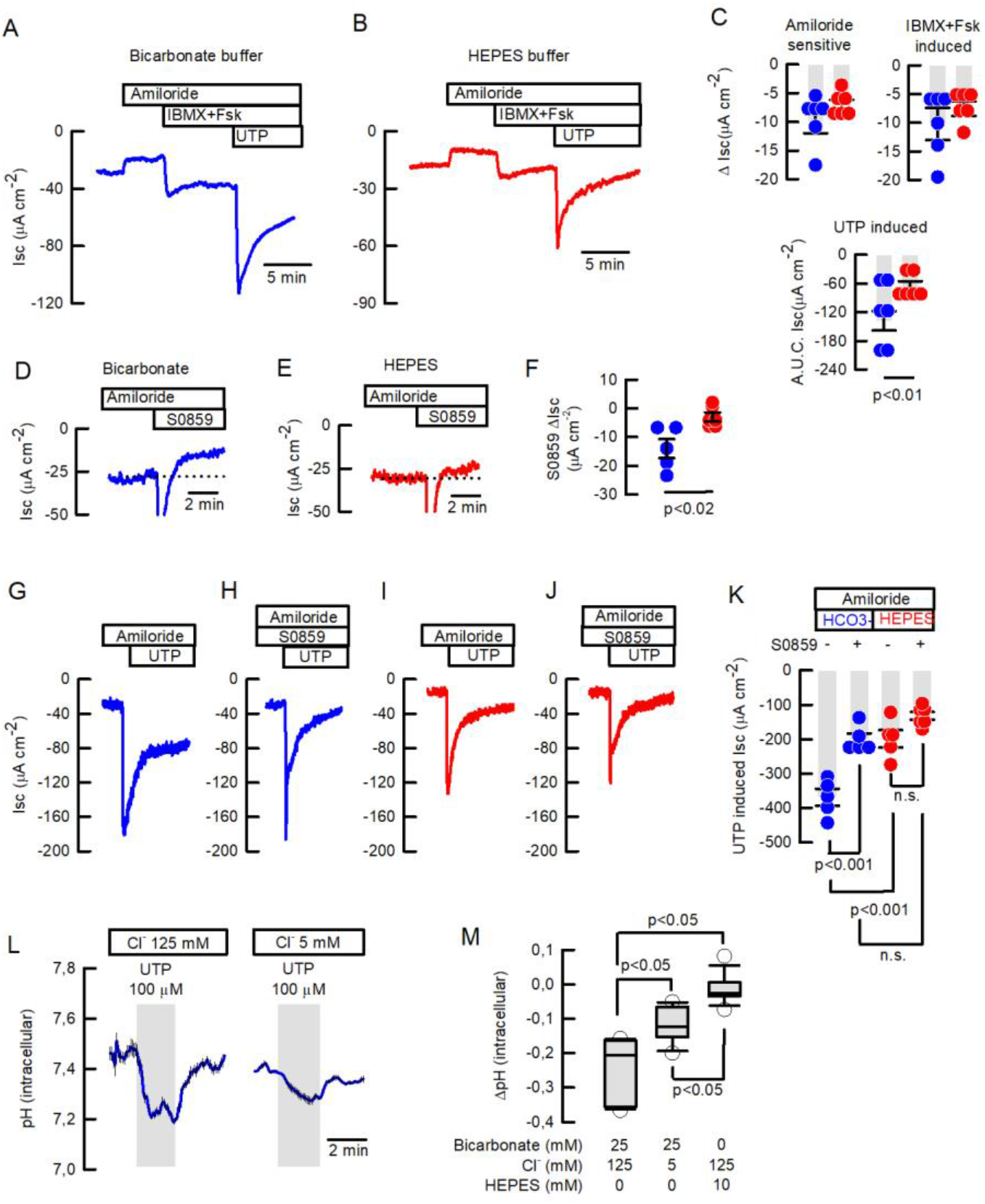
Basal and inducible bicarbonate secretion relies in SLC4A4 activity in mouse tracheal epithelium. Representative traces of I_SC_ in (A) bicarbonate and (B) HEPES buffer of freshly excised mouse tracheas. (C) Summary of ΔI_SC_ values for amiloride-sensitive Na^+^ absorption, IBMX+Fsk-induced and A.U.C. I_SC_ UTP-induced anion secretion in mouse tracheas; n=6 for each condition; Mann-Whitney Rank Sum Test. Representative I_SC_ traces of S0859 effect on unstimulated tracheas in (D) bicarbonate and (E) HEPES buffer. (F) Summary of ΔI_SC_ values for S0859 effect, n= 5 for each condition; Mann-Whitney Rank Sum Test. Representative I_SC_ traces for UTP-induced anion secretion in absence (G and I) or presence (H and J) of 30 μM S0859 in buffer bicarbonate (G and H) or HEPES (I and J). All in the presence of 10 μM amiloride. (K) Summary of experiments as presented in G-J; n=5 but HEPES+S0859 n=6; ANOVA on Ranks. Bars are mean ± S.E.M. (L) Average response of determination of pH_i_ in epithelial cells isolated from wild type mouse trachea and loaded with BCECF that were stimulated with 100 μM UTP and switched to 12.5 mM bicarbonate buffered solution. Experiments performed in bicarbonate buffer in blue and HEPES in red. (M) Summary of ΔpHi from experiments as those in (L) including a data set of cells maintained in HEPES buffer; Middle line of the box plot indicates the median; n=15 cells from 4 different mice, n=11 cells, 3 different mice and n=13 cells, 3 different mice respectively; ANOVA on Ranks.

Using the SLC4A4 blocker S0859, we observed the inhibition of the cAMP-induced anion current (ΔIsc -12.2 ± 2.4) suggesting that SLC4A4 might participate in the cAMP-response (Fig S3A-C). Nevertheless, when the TMEM16A/CFTR inhibitor, CaCCinhA01, was used to block the cAMP-induced current, further addition of S0859 was still able to induce a reduction in the current and of similar magnitude as shown in Fig S3A (Fig S3D-F; -10.1 ± 2.8 μA cm^-2^), confirming that electrogenic-bicarbonate secretion was not significantly stimulated by cAMP, indicating that basal HCO_3_^-^ secretion occurs in mouse trachea. To confirm this last hypothesis we added S0589 to tissues pre-incubated with amiloride and observed a reduction in the basal current only in HCO_3_^-^ buffer (−13.9 ± 3.3 vs -3.0 ± 1.6 Δ μA cm^-2^ for HCO_3_^-^ vs HEPES buffer; p<0.02; Mann-Whitney; Fig 3D-F). The magnitude of basal HCO_3_^-^ secretion inhibited by S0859 was similar to the experiments summarized in Fig S4C and 4F. Of note, the addition of S0859 to the tracheas induced a fast and transient negative change in I_sc_ as observed in Fig 3D and E and Fig S3A and D, that has been also observed in human cells (27), but for which the cause remains to be identified.

To further characterize the Ca^2+^-activated anion secretion, the UTP response was tested with no involvement of cAMP-induced secretion. As can be observed (Fig 3G,H) the UTP-induced anion secretion in tracheas maintained in HCO_3_^-^ buffer was significantly reduced by previous addition of S0859 (−368 ± 25 vs -200 ± 17 μA cm^-2^; p<0.001 One-way ANOVA). The UTP response was also reduced when HCO_3_^-^ was replaced with HEPES buffer (Fig 3I) (−199 ± 25 μA cm^-2^; p<0.001 One-way ANOVA), but the addition of S0859 induced no significant reduction of the UTP-induced anion secretion in tissues maintained in HEPES buffer (Fig 3J) (−142 ± 12 μA cm^-2^; p>0.05 One-way ANOVA).

### SLC4A4 participates in intracellular pH homeostasis in mouse airway epithelial cells

We reasoned that the UTP-induced HCO_3_^-^ exit would lead to cytoplasmic acidification and therefore we monitored intracellular pH of BCECF-loaded murine airway cells. As shown in Fig 3L, UTP induced an intracellular acidification (ΔpH -0.25 ± 0.02) that was significantly reduced when cells were placed in low Cl^-^ buffer (ΔpH -0.11 ± 0.02), indicating the existence of Cl^-^/HCO_3_^-^ exchange. Fig 3M summarizes changes in intracellular pH and includes experiments in HEPES buffer, which shows that UTP was almost unable to induce intracellular acidification (ΔpHi -0.01 ± 0.01) in absence of HCO_3_^-^. To test if the UTP-induced intracellular acidification was dependent on SLC4A4 activity, we tested the S0859 inhibitor and observed acidification of the intracellular compartment and prevention of UTP-induced acidification (Fig S3G-H). Washout of S0859 partially restored pHi and UTP-induced acidification. (Fig S3G-H; -0.05 ± 0.01 vs -0.13 ± 0.03 ΔpHi, for UTP with S0559 and UTP post washout respectively; p>0.002; Mann-Whitney). These data suggest that both basal and UTP-induced HCO_3_^-^ secretion are dependent on SLC4A4 activity in mouse airway epithelial cells.

### The genetic inactivation of *Slc4a4* induces a cystic fibrosis-like phenotype in mouse airways

As explained in the methods section, we decided to work with wild type and *Slc4a4*^-/-^ on the hybrid background, at 16-20 days of age. First, we observed that *Slc4a4*^-/-^ animals were affected by defects in tracheal cartilage formation with presence of ventral gaps and abnormal patterns on the rostrocaudal side (Fig 4A). In wild type animals, immunolocalization of SLC4A4 showed strong localization in the airway epithelium but the signal was nearly absent in the airways from the *Slc4a4*^-/-^ mice (Fig 4B). Using the same antibody, we showed that SLC4A4 was preferentially expressed in CCSP-positive cells that correspond to Club cells and was excluded from cells positive to acetylated-Tubulin, that identify ciliated cells (Fig 4C). This pattern of expression was maintained in distal airway bronchi and bronchioles (Fig S4B). Further histological examination of the *Slc4a4*^-/-^ mouse airways demonstrated the presence of adherent mucus at the surface of the tracheal epithelium (Fig 4D and S4C,D) as well as in the bronchi (Fig S4E-G) and bronchioli (Fig S5H). Signs of damaged epithelium was also observed as interruptions in the epithelial layer facing the lumen in the *Slc4a4*^-/-^ airways (Fig 4D and S4F,H), that might explain the decreased R_te_ of the tracheas placed in Ussing chambers and that prevented electrophysiological measurements in the *Slc4a4*^-/-^ tracheas (Fig 4E).

**FIGURE 4.**
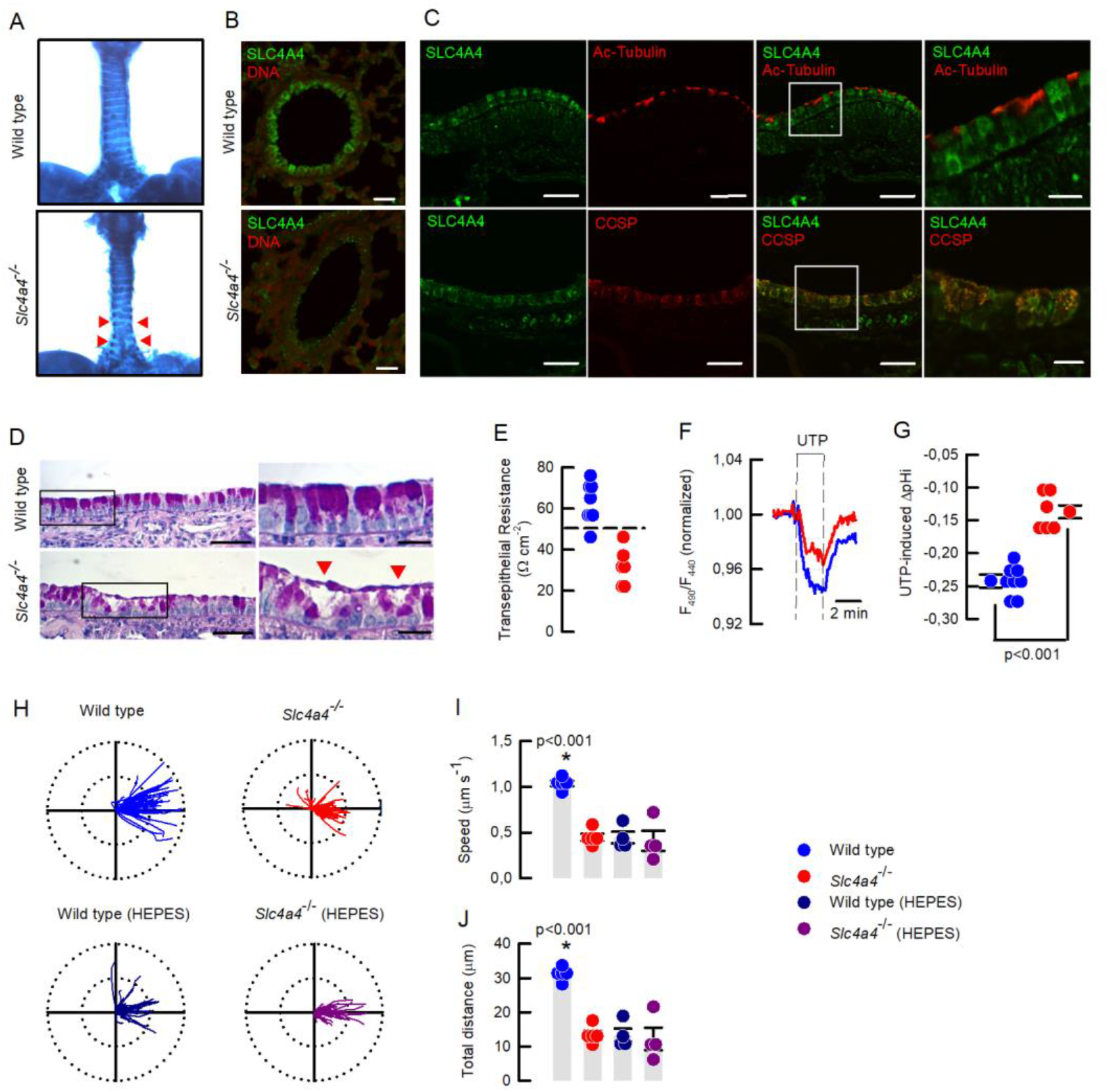
The *Slc4a4*^-/-^ bear signs of muco-obstructive airway disease. (A) Ventral view of Alcian blue stained tracheas of 17 days old mice. Red arrow heads show incomplete cartilage rings in the *Slc4a4*^-/-^ mouse; representative image of 3 animals per genotype. (B) SLC4A4 staining of epithelial cells is absent in the *Scl4a4*^-/-^ lung tissues; representative images of 3 different animals per genotype. (C) Epithelial SLC4A4 co-localizes with CCSP; representative images of 3 different animals. (D) Mucin staining showing mucus adhered to the epithelial surface and epithelial damage (red arrow heads); representative images of 5 animals per genotype. (E) Summary of R_te_values for wild type and *Scl4a4*^-/-^ tracheas; dashed line indicates 50 Ωcm^-2^; n= 8 for wild types and n=6 for *Scl4a4*^-/-^ tracheas. (F) Representative traces of pH_i_ in airway cells from wild type (blue) and *Slc4a4*^-/-^ (red) animals and (G) summary of UTP-inducedΔpH_i_ including mean ± S.E.M. and individual cells; n=8 cells for wild type and n=6 cells for *Scl4a4*^-/-^, 3 different animals per genotype; Mann-Whitney Rank Sum Test. (H) Beads tracking of MCC experiments for wild type and *Slc4a4*^-/-^ tracheas bathed with basolateral bicarbonate or HEPES buffer. Radius of the polar plots is 50 μm. Summary of MCC experiments for (I) speed, (J) total distance covered by beads. Bars indicate mean ± S.E.M.; n=5 for each genotype in bicarbonate buffer and n=4 for each genotype in HEPES buffer; ANOVA on ranks.

### Genetic silencing of *Slc4a4* impairs intracellular pH homeostasis and mucociliary clearance in mouse airways

In order to validate the role of SLC4A4 in pH homeostasis of murine airways, UTP-induced intracellular acidification was studied in airway cells isolated from the *Slc4a4*^-/-^ mice. We observed a decrease in the magnitude of intracellular acidification in *Slc4a4*^-/-^ cells when compared to those from wild type animals during UTP stimulation (−0.24 ± 0.01 vs -0.14 ± 0.01 ΔpH_[i]_, for wild type vs *Slc4a4*^-/-^) suggesting that an important amount of HCO_3_^-^ accumulates in airways cells via SLC4A4 (Fig 4F-G). We also noticed that after UTP wash-out, the acidification persisted in the wild type cells but not in the *Slc4a4*^-/-^ (Fig S4J; -0.09 ± 0.02 vs -0.02 ± 0.01 ΔpH_[i]_, for wild type vs *Slc4a4*^-/-^), suggesting that HCO_3_^-^ secretion was sustained by SLC4A4 activity. Clearance of plastic beads, as a way to measure MCC, was studied in freshly isolated mouse tracheas whose mucosal side was exposed to air. As shown in the polar plots in Fig 4H the plastic beads covered a larger distance in wild type tissues and, in some cases, retrograde movement of beads was observed in the *Slc4a4*^-/-^ trachea. A similar reduction in distance travelled was observed in wild type tissues bathed in HEPES buffer. The speed of plastic bead movement and total distance covered are summarized in Fig 4I,H. The use of HEPES buffer in the *Slc4a4*^-/-^ tracheas showed no further effect on both speed and total distance travelled by the beads. This data set demonstrates that mucociliary clearance is significantly decreased when HCO_3_^-^ transport is impaired after *Slc4a4* silencing.

## DISCUSSION

### SLC4A4 is a critical component of the bicarbonate secretory machinery

In this study we have established that the Na^+^-coupled HCO_3_^-^ transporter SLC4A4 or NBCe1 is central for bicarbonate transport, coupling to apical proteins to efficiently deliver bicarbonate in human and mouse airways. Early characterization in canine and human airway epithelium indicated that HCO_3_^-^ secretion occurs via CFTR, is activated by cAMP and dependent on basolateral Na^+^ (28), similar features were observed in the Calu-3 cell line too (29). Our experiments in hAECs corroborated the cAMP-activation and Na^+^-dependence, that has also been observed by others in the same (14, 27) or other cell types (30). Nevertheless, and in contrast to human airway epithelium, our present data shows that HCO_3_^-^ secretion in the mouse airways is mostly Ca^+2^-rather than cAMP-activated, but both human and mouse share a basal secretory component for HCO_3_^-^ previously observed in human bronchioles (31), and that we now show is dependent on SLC4A4 activity.

Basolateral localization of SLC4A4 was previously demonstrated in the Calu-3 cell line (32) but up to the date there has been no other studies investigating SLC4A4 localization in native airway tissues. Here we demonstrate that human airways express SLC4A4 in ciliated cells preferentially, and that the observed basolateral localization correlates with the functional evidence provided, reinforcing the role SLC4A4 in basolateral HCO_3_^-^ transport into the cells. Nevertheless, it must be considered that the sole expression of SLC4A4 does not assure HCO_3_^-^ secretion as compelling evidence has demonstrated that apical CFTR is also necessary. While the calcium-activated TMEM16A channel has been shown to be expressed in goblet cells (33), both HCO_3_^-^ transporters, CFTR and pendrin, have been localized in the apical membrane of ciliated hAECs cells supporting our observation (14, 34). Moreover, it has been demonstrated that SLC4A4-B, that we detected in human and mouse, and not the isoform A, is functionally coupled to CFTR through the IRBIT protein in pancreatic ducts (35, 36), further supporting the idea that a functional coupling in the same cell type is necessary for efficient HCO_3_^-^ secretion. CFTR distribution in airway epithelium has been under renewed scrutiny after the recent discovery of the pulmonary ionocyte, a rare airway epithelial cell type that contains the highest amount of *CFTR*/*Cftr* transcripts in human and mouse (37, 38). Nevertheless, recent and detailed studies demonstrate that most human airway epithelial cell types express CFTR including ciliated cells as was initially demonstrated (39, 40).

We noticed that SLC4A4 localization in mouse was different to that in human airway epithelium, as the mouse SLC4A4 was expressed in CCSP+ and not ciliated cells. Previously, CFTR and TMEM16A channels were specifically located in non-ciliated cells of mouse airways (41), a distribution also conserved in the rat (42). Pendrin, was also detected in non-ciliated secretory MUC5AC+ cells, suggesting that a functional coupling for HCO_3_^-^ secretion occurs in secretory non-ciliated cells of the mouse airways (43). Such a difference in expression of transporter proteins among human and mouse airways has frequently been described; for example CFTR, is not the principal Cl^-^ transporter in the mouse airways and consequently silencing or mutation of *Cftr* in mice did not produce CF disease of the lungs (44, 45). Furthermore, the expression in human and not mouse airways of another protein that influence ASL pH, the ATP12A H^+^/K^+^ ATPase, magnifies ASL acidification in human airways in CF disease (7).

### Impaired bicarbonate secretion produces a CF-like phenotype in the mouse airways

We demonstrated that SLC4A4 activity is pivotal for HCO_3_^-^ secretion. Inhibition of SLC4A4 induced acidification of the ASL in hAECs, an observation that suggests altering HCO_3_^-^ delivery can initiate a CF-like phenotype, and that was confirmed in the *Slc4a4*^*-/-*^ mouse model. Previous observations obtained by inducing silencing of *Tmem16a* in the mouse, or using cells carrying natural mutations in CFTR, affected both Cl^-^ and HCO_3_^-^ secretion making it difficult to understand what the consequences of reduced transport for each anion alone are (27, 46). As observed here, the silencing of *Slcl4a4* induced a muco-obstructive phenotype in the mouse, whereas the silencing of *Slc12a2*, which encodes the bumetanide-sensitive NKCC1 co-transporter, which is essential for Cl^-^ accumulation and secretion, did not. This suggests that lack of HCO_3_^-^ is pathologically more relevant than Cl^-^ secretion in mouse airways (47). Indeed, a significant amount of experimental evidence, using different models and species, is consistent with our findings in the *Slc4a4*^-/-^ mouse. For example, the acidification of the ASL induced abnormal epithelial immune responses, increased mucus viscosity and reduced MCC that could be reversed after HCO_3_^-^ supplementation (6, 7, 48–51). Furthermore, HCO_3_^-^ secretion is necessary for proper mucus release from hAECs (27), and maintenance of normal amounts of ENaC-mediated Na^+^ absorption and ASL volume (8), functions that become abnormal due to acidic ASL pH. Even though technically challenging issues prevented us from measuring ASL pH in the mouse trachea, the reduction of HCO_3_^-^ transport demonstrated here after SLC4A4 inhibition is consistent with the expected muco-obstructive phenotype, including reduced MCC and mucus accumulation.

### Extra-renal phenotypes after *Slc4a4* inactivation and human mutations

Mouse models of *Slc4a4* silencing have shown differences in phenotypes depending on the isoforms affected. The *Slc4a4*-null animal, corresponding to the one used in the present studies, and the *Slc4a4*^W516X/W516X^ avatar mouse engineered to mimic a non-sense mutation found in a human patient, are affected by severe metabolic acidosis due to proximal renal tubular acidosis (pRTA)(21, 52). Even though, our results showing mucus accumulation in the *Slc4a4*^-/-^ mouse airways might be influenced by the decreased availability of plasma HCO_3_^-^ in the model tested (5.3 ± 0.5 mM)(21), we clearly demonstrated that SLC4A4 activity is necessary for maintenance of ASL pH and airway clearance discarding such a possibility.

Extra-renal manifestations of *Slc4a4* silencing in the mouse include growth retardation, ocular band keratopathy, splenomegaly, abnormal dentition and intestinal obstructions, most of which mimics the clinical findings observed in human patients. Nevertheless lung defects have not been reported in human patients which could be explained by the fact that human disease is milder than in mouse models. This might be related to the fact that at birth human kidneys are functionally more mature than in mouse (53) and for example, while heterozygous animals present mild pRTA, human patients bearing heterozygous SLC4A4 mutations did not show any signs of disease (21, 52). Even though, a compensatory activity of other Na^+^-coupled HCO_3_^-^ cotransporters, or exchangers, has been discarded in mouse, this possibility has not been examined in humans where compensatory activity might benefit patient’s health (52).

Both the *Slc4a4*^-/-^ and W516X-mutant animals die soon after weaning as we also observed, but while the specific knock-out mouse for SLC4A4 isoform A (NBCe1-A) has a normal life span, the knock-out mouse for SLC4A4 isoforms B and C (NBCe1-B/C) is still lethal. This suggests that the cause of death of the animals was not due to metabolic acidosis, but rather due to extra renal phenotypes worsened by pRTA (54, 55). This observation might also be linked to the influence of modifier genes as observed in human patients and CF mouse models (56–58). In this regard, our observation that lethality is dependent on the genetic background of the animals supports such a possibility.

It will be of interest to use the *Slc4a4* animal models and generate an airway specific null mouse to study the pathogenesis of muco-obstructive diseases of the lungs due to reduced HCO_3_^-^ secretion. We believe that the elucidation of the transport systems that participate in pH maintenance in the airways offers the chance of increasing our current knowledge of the impact of impaired bicarbonate transport during health and disease.

## ACKNOWLEDGMENTS

This work was supported by two CF Trust Strategic Research Centre grants (SRC003 and SRC013) and a Medical Research Council (MRC) Confidence in Concept grant (MC_PC_15030). The Centro de Estudios Científicos (CECs) is funded by the Base Financing Programme of CONICYT, Chile. Cells from Dr. Randell were supported by Cystic Fibrosis Foundation grant (BOUCHE15R0) and NIH grant (P30DK065988). We would like to acknowledge Drs. Scott H. Randell and Leslie Fulcher (Marsico Lung Institute, The University of North Carolina at Chapel Hill, United States) for providing human primary airway epithelial cells from the UNC CF Center Tissue Procurement and Cell Culture Core. Git Chung for human kidney RNA sample.

---

**Supplemental Figure 1.**
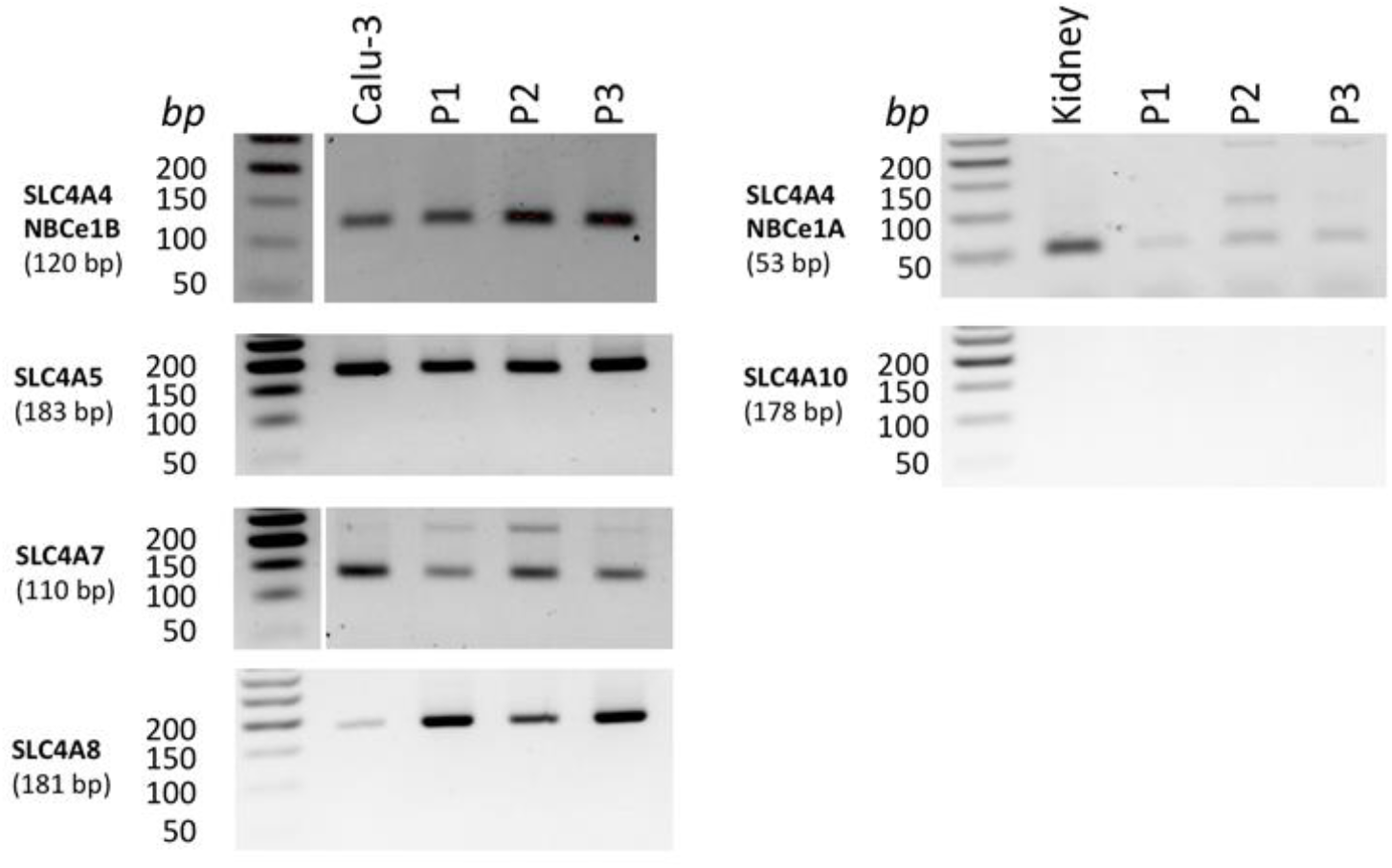
Expression of SLC4A family members of bicarbonate transporters in differentiated primary human airway epithelial cells. Amplicons for *SLC4A4, SLC4A5, SLC4A7, SLC4A8* and *SLC4A10* were amplified from differentiated human airway epithelial cells from 3 donors. Pancreatic and renal isoforms of SLC4A4 were detected in the airway epithelial cell line Calu-3 (lane 2, left panels) and human kidney (lane2, right panels).

**Supplemental Figure 2.**
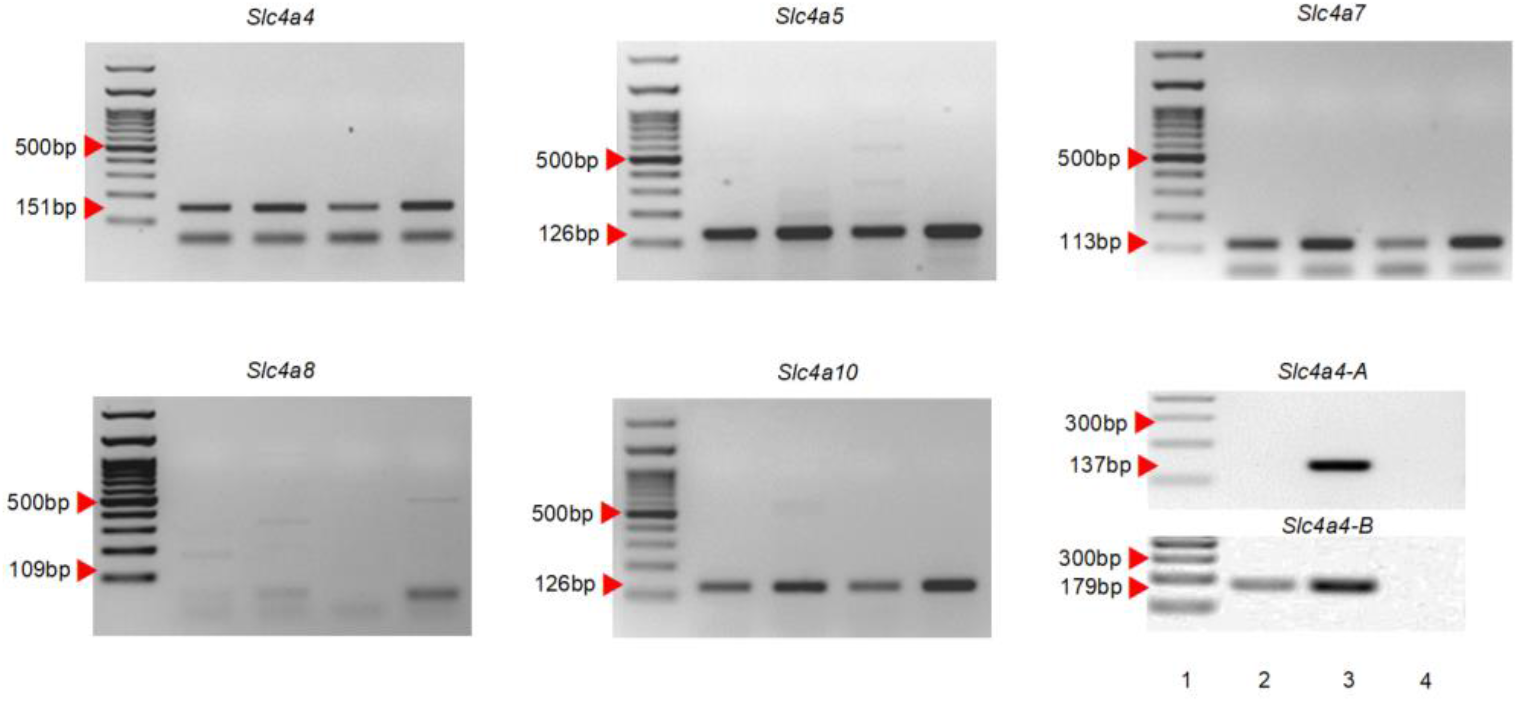
Expression of SLC4 family members of bicarbonate transporters in mouse airway epithelial cells. Amplicons for *Slc4a4, Slc4a5, Slc4a7, Slc4a8* and *Slc4a10* were amplified from 4 samples of airway epithelial cells isolated from 4 different animals. Isoforms of *Slc4a4* were detected in airway epithelial cells (lane 2) and mouse kidney (lane3). Lane 4 corresponds to negative RT-PCR control. Image representative of 4 samples tested.

**Supplemental Figure 3.**
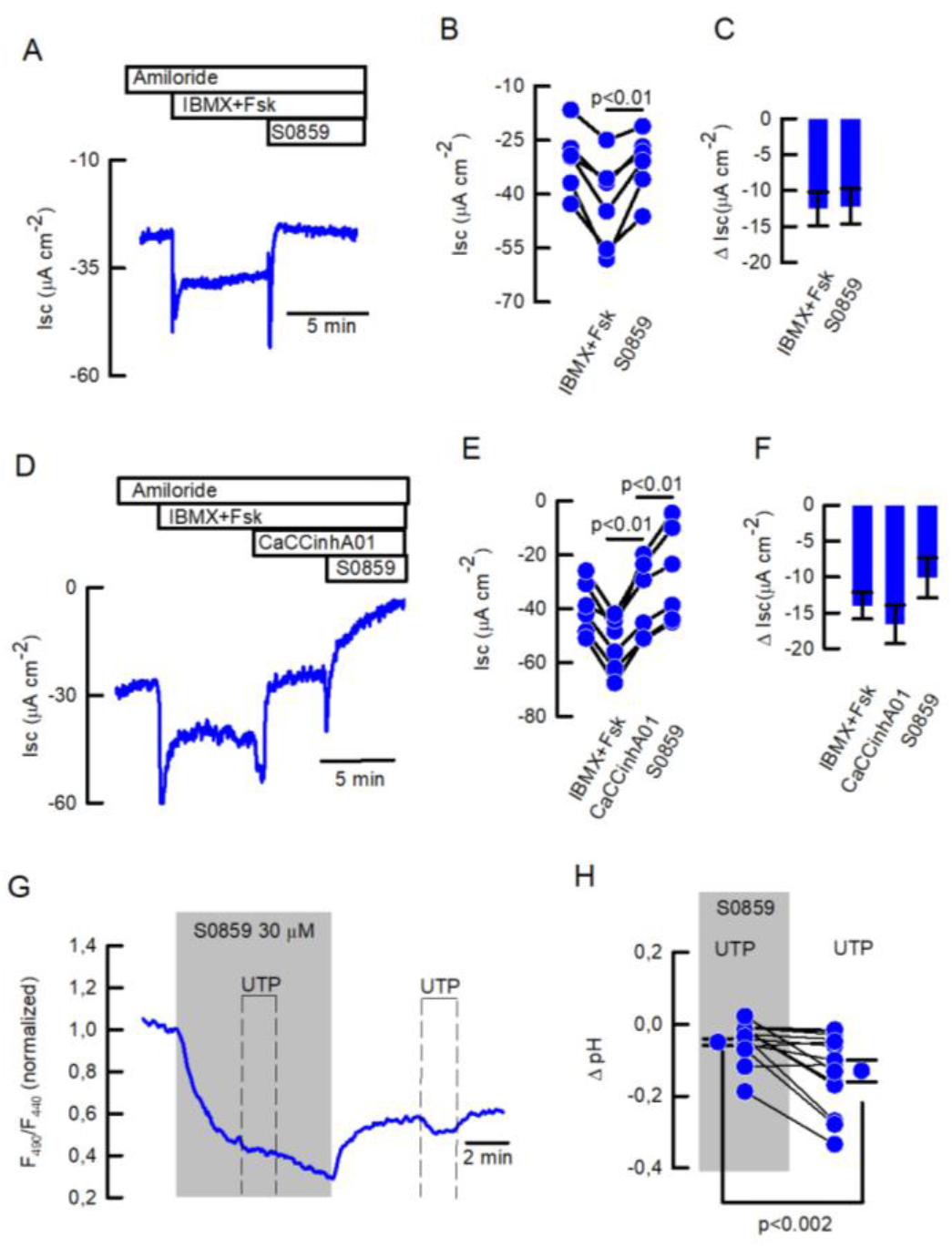
(A) Representative trace of S0859 effect on IBMX+Fsk-induced anion secretion, (B) individual experiments showing Isc changes and (C) summary of ΔI_SC_ values for IBMX+Fsk-induced anion secretion and S0859. (D) Representative trace of S0859 (30) effect on IBMX+Fsk-induced anions secretion after inhibition of apical chloride secretion by addition of CaCCinhA01 (30 μM), (E) individual experiments showing I_SC_ changes and (F) summary of ΔI_SC_ values for experiments in (D). (H) UTP-induced intracellular acidification in the presence of 30 μM S0859 in bicarbonate buffer or HEPES. (I) Summary of experiments showing ΔpH_[i]_; n=14 cells from 4 different mice; Mann-Whitney Rank Sum Test.

**Supplemental Figure 4.**
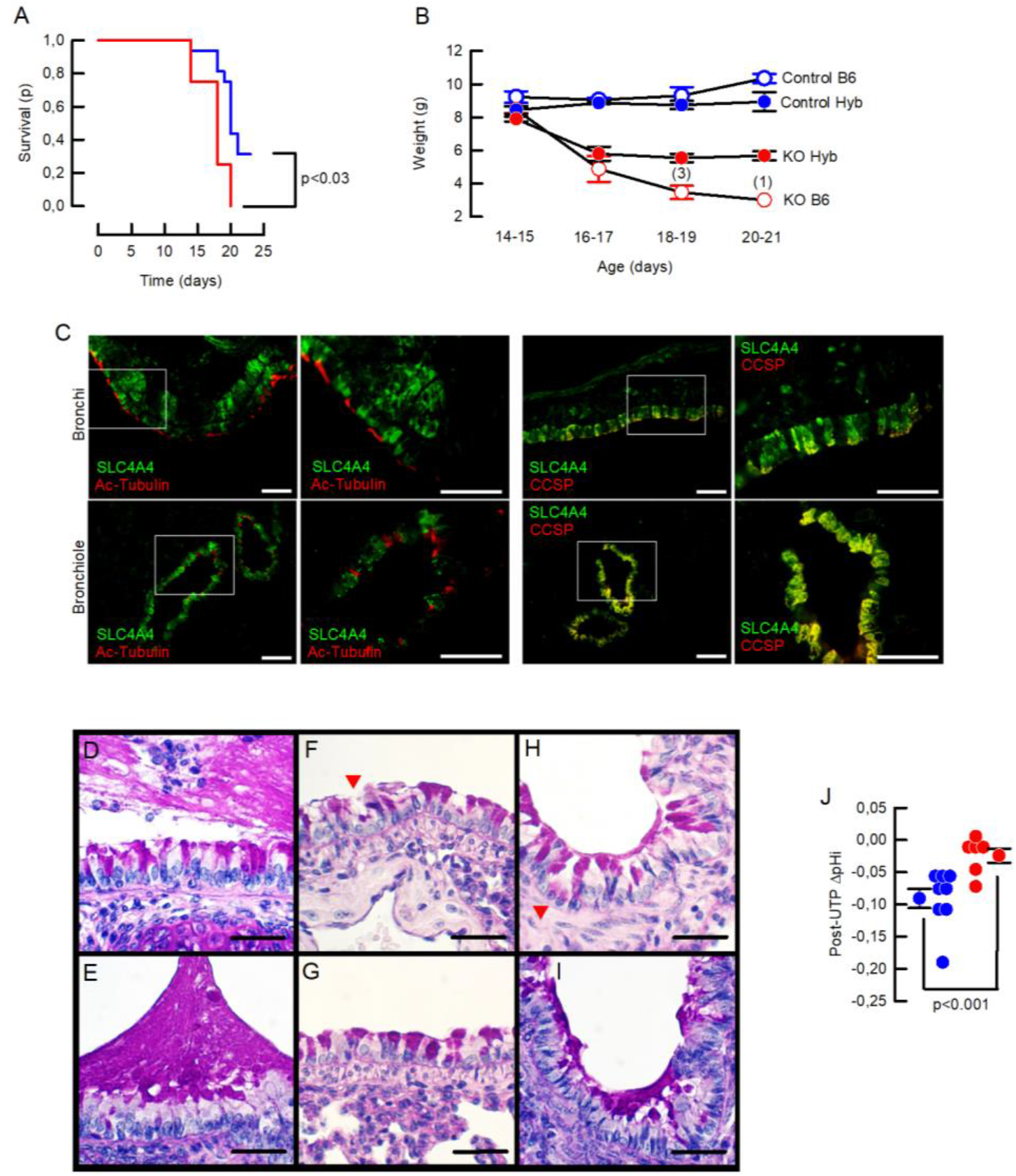
*Slc4a4* silencing produces early lethality, weight loss, mucus accumulation and reduced intracellular pH. (A) Kaplan-Meier curves showing survival for *Slc4a4*^-/-^ in the C57Bl6/J (B6; red line) or hybrid background (Hyb; blue line). Log-Rank test; n=12 and 16 per gruop. (B) Weight gain curves for control and *Slc4a4*^-/-^ (KO) mice in B6 and Hyb backgrounds; n=5 for each group but for KO B6 that due to increased lethality group is reduced as indicated in the numbers on top of last 2 records. (C) Epithelial SLC4A4 co-localizes with CCSP protein in bronchi and bronchioles; representative images of 3 animals per genotype. (D-I) PAS staining of mucus in trachea (D-E), bronchi (F-H) and bronchiole (I). Red arrows indicate areas of epithelial damage. Images are representative of 5 different animals per genotype. (J) Summary of ΔpH_[i]_ post-UTP including mean ± S.E.M. and individual cells; n=8 cells for wild type and n=6 cells for *Scl4a4*^-/-^, 3 different animals per genotype; Mann-Whitney Rank Sum Test.

**Suplemetal Table 1.**
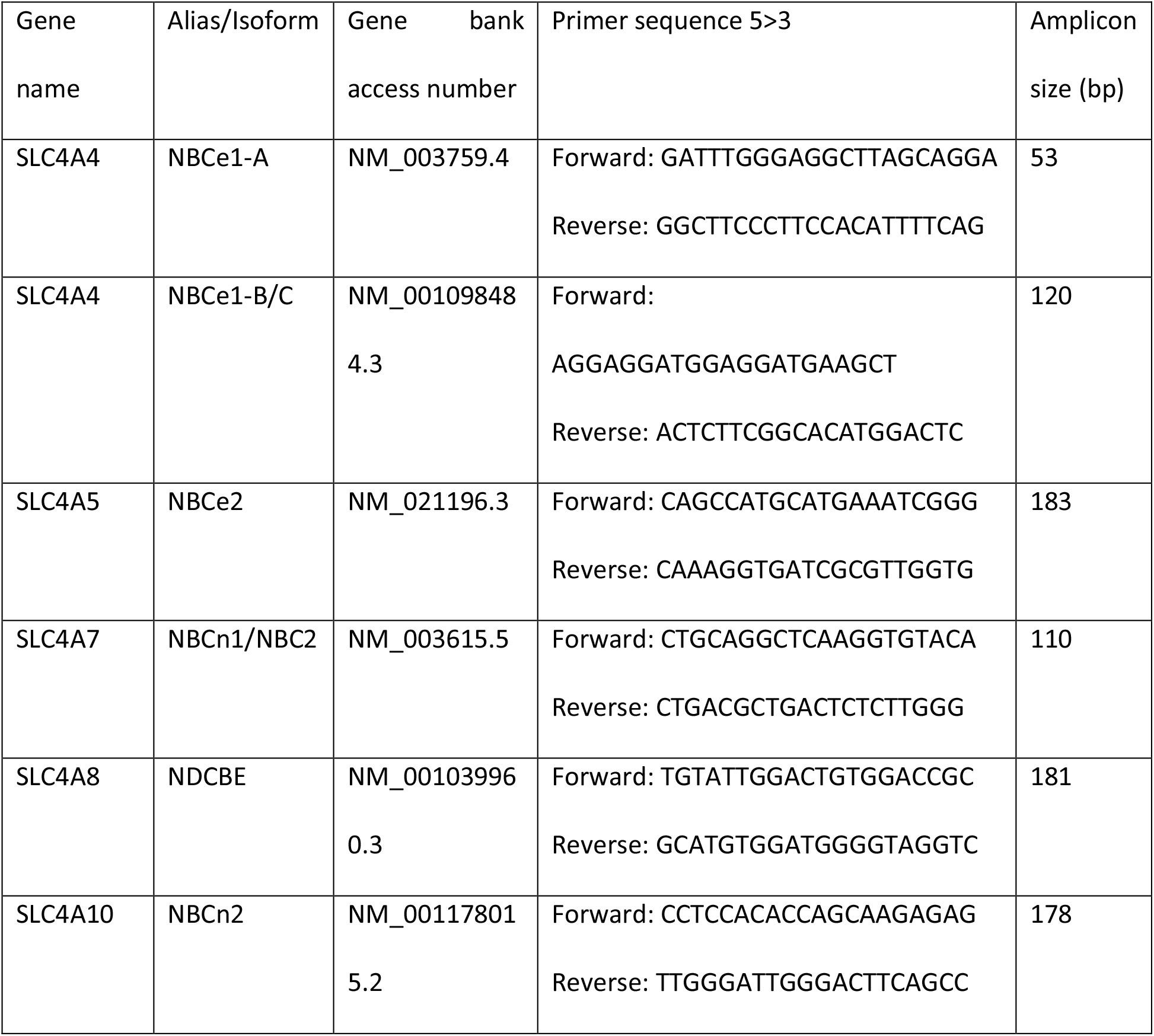
Primers for hBECs.

**Suplemetal Table 2.**
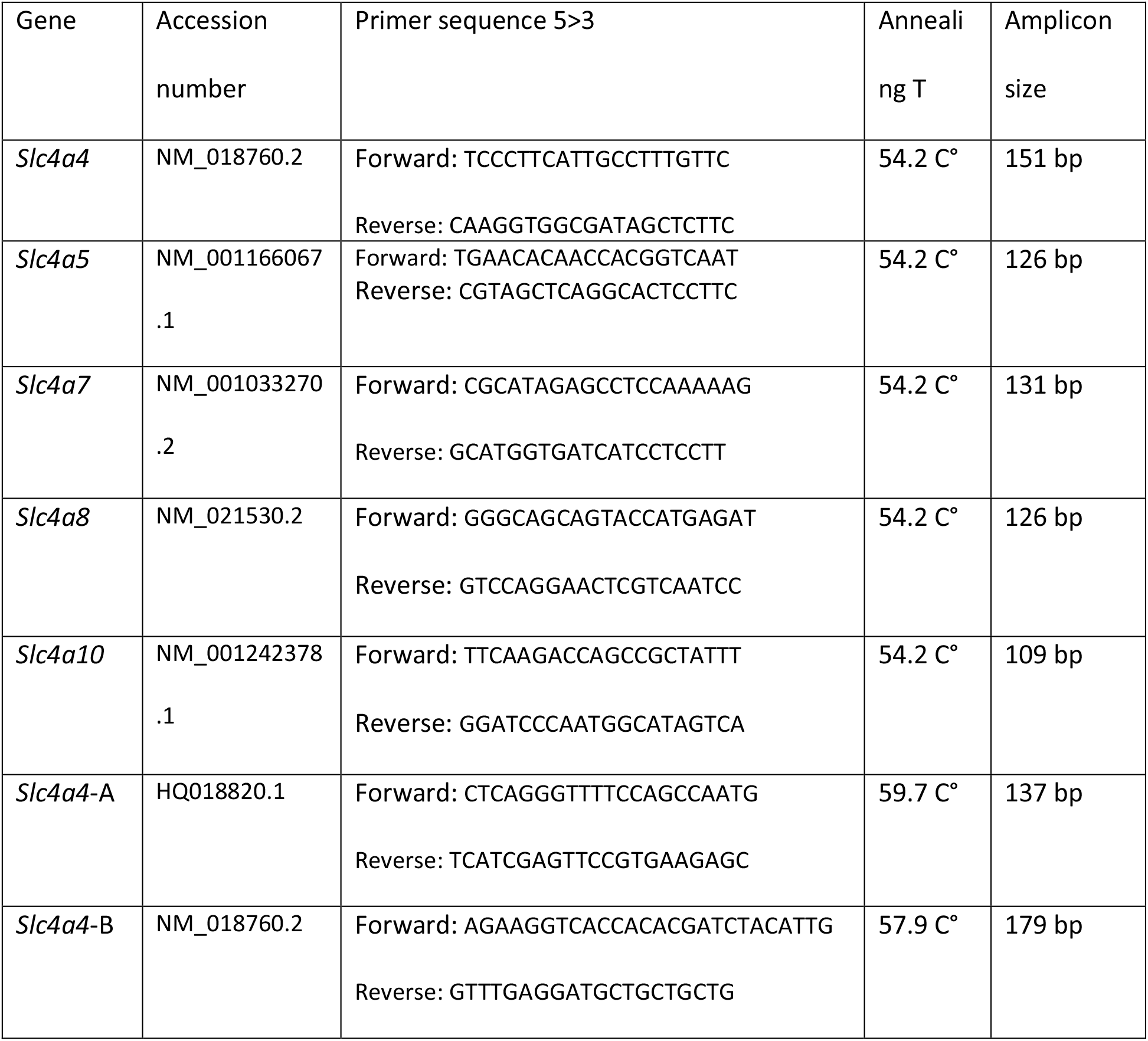
Primers for mouse.

